# Specific and non-uniform brain states during cold perception in mice

**DOI:** 10.1101/2022.10.20.513008

**Authors:** Haritha Koorliyil, Jacobo Sitt, Isabelle Rivals, Yushan Liu, Silvia Cazzanelli, Adrien Bertolo, Alexandre Dizeux, Thomas Deffieux, Mickael Tanter, Sophie Pezet

**Author notes:** Corresponding Author: Sophie PEZET, Address: Institute of Physics for Medicine Paris, ESPCI Paris, 17 rue Moreau, 75012, Paris, France, <u>Email</u>.

## Abstract

The quest to decode the complex supraspinal mechanisms that integrate cutaneous thermal information in the central system is still ongoing. The dorsal horn of the spinal cord is the first hub that encodes thermal input which is then transmitted to brain regions via the spinothalamic and thalamo-cortical pathways. So far, our knowledge about the strength of the interplay between the brain regions during thermal processing is limited. To address this question, we imaged the brains of awake and freely-moving mice using Functional Ultrasound imaging during plantar exposure to constant and varying temperatures. Our study, a synchronous large field investigation of mice brains reveals for the first time the brain states and the specific dynamic interplay between key regions involved in thermal processing. Our study reveals: i) a dichotomy in the response of the somato-motor-cingulate cortices and the hypothalamus, which was never described before, due to the lack of appropriate tools to study such regions with both good spatial and temporal resolutions. ii) We infer that cingulate areas may be involved in the affective responses to temperature changes. iii) Colder temperatures (ramped down) reinforces the disconnection between the somato-motor-cingulate and hypothalamus networks. iv) Finally, we also confirm the existence in the mouse brain of a dynamic brain mode characterized by low cognitive strength, described previously only in non-human primates and humans. The present study points towards the existence of a common hub between somato-motor and cingulate regions, whereas hypothalamus functions are related to a secondary network.

## INTRODUCTION

Thermal sensation and perception are crucial for maintaining the structural and functional integrity of all organisms (Gracheva and Bagriantsev, 2015). Thermal changes can elicit a multitude of responses including rapid motor withdrawal reflex and thermoregulation to maintain core body temperature. To cope with the changes in the thermal environment, physiological and behavioral mechanisms are employed permanently (Tan and Knight, 2018). The complex mechanisms that result in the central and peripheral integration of cutaneous thermal sensations is still not completely understood.

Thermal sensations felt on skin are encoded and transmitted to the central nervous system by primary sensory neurons, such as non-myelinated C fibers and thinly myelinated Aδ fibers whose terminals act as free nerve endings on the skin (Middleton et al., 2021; Xiao and Xu, 2021). The thermal information is transmitted by the sensory afferents to the dorsal horn via the TRP channels, which are the molecular thermo-detectors (Peier et al., 2002; Clapham, 2003; Patapoutian et al., 2003; Bandell et al., 2004; Bautista et al., 2007; Dhaka et al., 2007; Tan and McNaughton, 2016; Hoffstaetter et al., 2018; Vandewauw et al., 2018; Vilar et al., 2020), and can initiate well-defined responsive pathways, such as: a) activation of motor neurons resulting in a rapid withdrawal reflex, b) transmission of thermal information via the spinothalamic tract to various nuclei of the thalamus and finally to several cortical areas such as the insular cortex and the primary somatosensory cortex where it takes the form of a perceived temperature, c) initiation of thermoregulatory responses (Vriens et al., 2014). Although the supraspinal pathways work synergistically to form a thermal perception, the complex interplay among them is less explored. Studies have shown that several thalamic nuclei (Bushnell et al., 1993; Duncan et al., 1993; Craig et al., 1994; Davis et al., 1999), somatosensory regions (Becerra et al., 1999; Moulton et al., 2012; Milenkovic et al., 2014) and the insula (Craig et al., 2000; Olausson et al., 2005; Veldhuijzen et al., 2010; Peltz et al., 2011; Wager et al., 2013; Gogolla et al., 2014) are crucial for thermosensation. Although the cingulate region is not a direct part of the thermosensory circuit, it is involved in the affective responses to the nociceptive thermal stimulations (Vogt, 2005). The preoptic anterior hypothalamus (POAH) has been linked to thermoregulatory behavior in numerous studies (Ishiwata et al., 2002; DiMicco and Zaretsky, 2007; Wang et al., 2019).

The present study aimed at understanding the involvement of some of the aforementioned brain regions using functional ultrasound (fUS) imaging, which is a relatively new versatile neuroimaging modality that allows imaging and measurement of cerebral blood volume in humans (Demene et al., 2017; Imbault et al., 2017; Soloukey et al., 2020), non-human primates (Dizeux et al., 2019) and rodents (Macé et al., 2011; Sieu et al., 2015; Urban et al., 2015; Bergel et al., 2018; Rahal et al., 2020) with excellent spatial (100 to 300 μm) and temporal resolutions (down to 20 ms). One of its most important characteristics is its high sensitivity compared to fMRI (Boido et al., 2019). Indeed, during a task, due to neurovascular coupling, the locally increased neuronal activity leads to a strong hemodynamic response (Iadecola, 2017). In the past, fUS imaging proved sensitive enough to enable the measurement of the cortical hemodynamic changes induced by sensory (Macé et al., 2011), olfactory (Osmanski et al., 2014a) and visual (Macé et al., 2018) stimuli in anesthetized animals. Another very important characteristic of fUS imaging consists in its ability to perform acquisitions in awake and behaving animals (Montaldo et al., 2022), as demonstrated for auditory stimuli in awake animals (Bimbard et al., 2018) or motor tasks (Sieu et al., 2015; Bergel et al., 2020). Taking advantage of the sensitivity of this technique and the ability to study freely moving animal, this study aimed at improving our understanding of the processing of warm and cold sensing by studying the changes of intrinsic brain connectivity in freely moving mice during plantar exposure to warm, cold and neutral surfaces, and to the variations of the surface temperature. The analysis of the static and dynamic functional connectivity reveals that cold induces a strongly increased connectivity in the somato-motor (SM) network, but also a decreased connectivity between the SM areas and the hypothalamus.

## MATERIALS AND METHODS

### ANIMALS

The experiments were conducted in compliance with the European Community Council Directive of 22 September 2010 (010/63/UE) and the local ethics committee (Comité d’ethique en matière d’expérimentation animale N° 59, “Paris Centre et Sud”, project 2018-05). Accordingly, the number of animals in our study was kept to the minimum necessary. Due to previous studies using a similar experimental design (Rabut et al., 2020), we established that N=6 animals per group was the smallest number of animals required to detect statistically significant differences in our imaging experiments. Finally, all methods are in accordance with ARRIVE guidelines.

Animals arrived at the animal facilities one week before the beginning of experiments. Twelve C57Bl/6 male mice (aged 7 weeks at the beginning of the experiments) were obtained from Janvier labs (France) and housed under controlled temperature (22 ± 1°C), relative humidity (55 ± 10%), with a 12-hour light/dark cycle. Finally, food and water were available ad libitum. Experiments typically lasted for 4-6 weeks.

Constant temperature and temperature ramp experiments were conducted in two different sets of N=8 and N=6 mice (Figure 1). When possible, animals were imaged more than once in each experimental condition. Indeed, due to motion artifacts, some sessions had to be discarded (see below). Details of data included from the various animals are listed in supplementary table 1.

**Figure 1:**
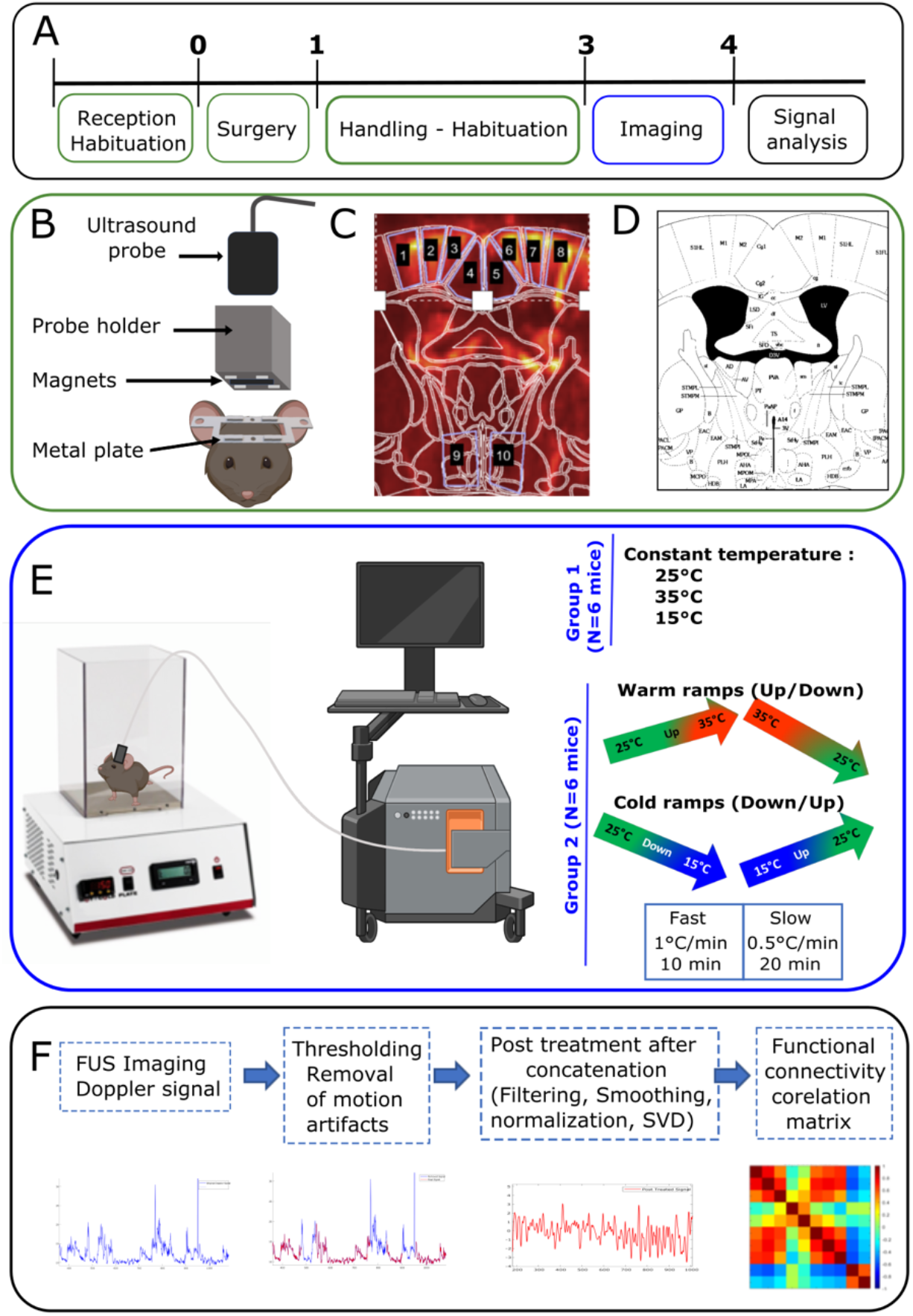
Experimental design and timeline of the experiments. A) One week after their arrival in the laboratory, a metal plate was surgically attached to the mice skulls. After 3 days of recovery, mice were handled and habituated for 2 weeks. Imaging was then performed for the next 2 weeks or more, depending on the quality of the skull and on the well-being of the mice. (B) Schematic showing the metal plate, probe holder, fUS probe and (C-D) plane of imaging (Bregma −0.34 mm). C shows the Doppler image superimposed with the delimitation of the mouse brain atlas (D, Paxinos and Franklin, 2011). The regions of interest are: 1,8 primary somatosensory cortex, hindlimb part, 2,3,7,8 primary and secondary motor cortices, 4,5 Cingulate cortex and 9,10 Hypothalamus. (E). Experimental setup with Bioseb and fUS Imaging (Iconeus). Mice were subjected to constant temperatures, or to warm and cold ramps at fast and slow pace. (F) The doppler signal obtained from imaging underwent thresholding to remove motion artefacts, concatenation, low-pass filtering, smoothing, normalization and SVD filtering. The cleaned doppler signal was then used for FC analysis.

### SURGICAL IMPLANTATION OF METAL PLATE

Approximately one week after their arrival, the mice underwent surgery for the implantation of the metal plate (Tiran et al., 2017; Rabut et al., 2020). A mixture of ketamine (100 mg/kg) and medetomidine (1 mg/kg) was administered intraperitoneally and then the mouse was placed on a stereotaxic frame where the skull bone was exposed after skin and periosteum removal. The metal plate was fixed on the skull using Superbond C&B (Sun Medical, USA) and small screws minimally drilled in the skull. The field of interest was approximately 5mm wide, between the Bregma and Lambda points. The surgery took 45 to 60 minutes to be completed. Subcutaneous injections of atipamezole (1 mg/kg, Antisedan) and Metacam (5 mg/kg/day) were given to reverse the anesthesia and to prevent postsurgical pain, respectively. A protective cap was mounted on the metal plate using magnets to protect the skull and to keep the field of imaging intact for 4-6 weeks (Bertolo et al., 2021). Altogether, the metal plate and the cap did not interfere with the normal daily activity of the mice. After a recovery period of 3 days, the mice proceeded towards the habituation phase.

### HABITUATION AND TRAINING

After recovering from the metal plate implantation, the mice were subjected to an extensive habituation protocol. To make sure that the mice were not under any stress during the experiment, it was crucial that they were at ease with the user and the setup. They were initially handled by the user and then exposed to the Bioseb HC Plate. Their daily interaction time in the Bioseb apparatus was gradually increased from 15 minutes to 30 minutes, 1 hour, 1 hour 30 minutes and finally 2 hours. Depending on the level of habituation of each mouse, the user practiced the protective cap removal, the skull cleaning using saline and application of echographic gel without anesthesia, by gently restricting the head movement. After each session, the mice received a reward. The process lasted 2 weeks and, depending upon the comfort level of the mice, we then proceeded to the imaging phase.

### EXPERIMENTAL PARADIGM

The global aim of this study was to decipher how thermal sensations are encoded in the mouse brain. As intrinsic functional connectivity was shown to measure the activity and functionality of the brain networks, we postulated it could vary during exposure to cold, warm and neutral floor surfaces. To address this issue, we imaged the brain using fUS in awake and freely moving mice exposed to various thermal sensations (Figure 1 E).

We observed in previous experiments (Rabut et al., 2020) that motion artifacts deeply alter the quality of the ultrasound signals, and that all efforts need to be made to prevent these artifacts. In preliminary experiments, we have sought to establish the range of temperature in which the animals were minimally uncomfortable. Combining the records of the animal’s natural behavior (grooming, exploration, urination and freezing) and measurements of naturally emitted ultrasound vocalizations, we observed that, between 15°C and 35°C, the animals were minimally uncomfortable and the experiments feasible. Any residual of motion artifacts in the Doppler signal (due to head movements or behavioral movements such as grooming or licking) were removed using a dedicated signal processing as explained in Figure 1F.

We used the Bioseb ‘Hot-Cold Plate’ to conduct the experiments. The metal floor of the Bioseb equipment can be kept at a constant temperature, or heated or cooled down at different rates.

#### i. Constant Temperature

A constant floor temperature of 15°C (cold), 25°C (neutral) or 35°C (warm) was applied for 20 minutes. Experiments at these temperatures were randomly repeated on two separate days. At the beginning of all sessions, the floor temperature was held at 25°C, and the transition to the desired temperature occurred within seconds.

#### ii. Varying Temperature at fast or slow pace

Floor temperature variations into the warm domain were made of two ramps, one up (from 25°C to 35°C) and one down (back to 25°C); variations into the cold domain were made of a ramp down (from 25°C to 15 °C) and a ramp up (back to 25°C) (Figure1 E). In order to determine the effect of the speed of the temperature change, these ramps were performed at 2 different rates: either at 0.5°C per minute (for 20 minutes) or at 1°C per minute (for 10 minutes). The mice were reimaged at least 3 times. In order to avoid any bias due to the order of these variations, the order of warm and cold variations was randomized. They are denoted as follows:

##### Cool Ramps

■ **CFD**: Cool Fast Down: 25°C to 15 °C (−1°C per minute)
■ **CFU**: Cool Fast Up: 15°C to 25° (+ 1°C per minute)
■ **CSD**: Cool Slow Down: 25°C to 15 °C (+ 0.5°C per minute)
■ **CSU**: Cool Slow Up: 15°C to 25°C (−0.5°C per minute)

##### Warm Ramps

■ **WFU**: Warm Fast Up: 25°C to 35°C (+1°C per minute)
■ **WFD**: Warm Fast Down: 35°C to 25°C (−1°C per minute)
■ **WSU**: Warm Slow Up: 25°C to 35°C (+0.5°C per minute)
■ **WSD**: Warm Slow Down: 35°C to 25°C (−0.5°C per minute)

### TRANSCRANIAL AWAKE FUS IMAGING

Three days prior to the first imaging session, the mice were anesthetized with isoflurane (1.5%). The respective probe holders were magnetically clipped to the metal plate. Real time transcranial Doppler images were acquired using the NeuroScan acquisition software (Inserm Technological Research Accelerator and Iconeus, Paris France), and the position of the probe was adjusted to select the Bregma −0.34 mm plane. The regions of interest included the primary somatosensory cortex of the hindlimb, the primary and the secondary motor cortex, the cingulate cortex and the hypothalamus (Figure 1 C-D). The skull was then thoroughly inspected and cleaned to avoid any infection. The mice were put back in their cages and imaged only 3 days later to avoid any interference with the isoflurane anesthesia. Unlike the previous awake imaging protocols described in (Tiran et al., 2017), mice were not anesthetized to prepare the skull during the imaging phase. They were trained and accustomed to the detachment of the protective cap, to the cleaning of the skull with saline, and to the application of echographic gel with minimal force. They were then gently introduced into the Bioseb apparatus and the probe holder was attached to the implanted metal frame using the magnets on both pieces. Experiments began shortly after.

Real time vascular images were obtained by ultrafast compound doppler imaging technique (Deffieux et al., 2021). Eleven successive tilted plane waves (−10° to +10° with 2° steps) were used for insonification. Each image was obtained from 200 compounded frames acquired at 500Hz frame rate corresponding to a 5.5 kHz pulse repetition frequency. The tissue signal was isolated from the cerebral blood volume signal using a spatio-temporal clutter filter based on the singular value decomposition (SVD) of raw ultrasonic data (Demene et al., 2015) to obtain a power Doppler image.

### DOPPLER SIGNAL ANALYSIS

Imaging in awake mice required careful removal of motion artifacts due to head movements or behavioral movements such as grooming. We followed the analysis previously described in (Rabut et al., 2020) by first using a SVD clutter filter to separate blood motion from tissue motion, and then by thresholding tissue motion and Doppler signal to identify the frames with motion artifacts. Several thresholds were investigated by carefully examining the tissue motion signal and the threshold that removed most of the motion artifacts was chosen. We kept and concatenated epochs of at least 50 consecutive time points. The concatenated cleaned frames were filtered using a low-pass filter with a cut-off frequency of 0.1 Hz to extract the steady-state. A polynomial fit of order 3 was applied to detrend the signal and, finally, global variations in the brain were suppressed by removing the first eigenvector of the dataset before connectivity analysis (Figure 1F). Acquisitions that did not match the aforementioned criteria (<50 consecutive timepoints) were discarded. Supplementary Table 1 summarizes the identify of animals included in both parts of the study, and how many sessions from each mouse were kept. Out of the N=8 animals included in the constant experiments, N=8 acquisitions per experimental conditions were used in the analysis. As for the second part of the study (ramp experiments), they included 6 to 10 acquisitions (Supplementary Table 1), obtained in N=6 mice.

### STATISTICAL ANALYSES

In order to understand the interactions between brain regions involved in the temperature coding, we performed static and dynamic FC analyses. In the two types of analyses, the following brain regions of the imaging plane were studied: Primary Somatosensory Hind Limb part, Primary and Secondary Motor, Cingulate and Hypothalamus (Figure 1 C, D). This imaging plane was chosen because of the known role of some of these regions in thermosensation and thermoregulation. Ten ROIs were defined based on the Paxinos Atlas (Paxinos, G and Franklin, 2012), numbered and coded as:

- 1 (S1HLL) & 8 (S1HLR): primary somatosensory cortex left & right,
- 2 (M1L) & 7 (M1R): primary motor cortex left & right,
- 3 (M2L) & 6 (M2R): secondary motor cortex left & right,
- 4 (CgL) & 5 (CgR): cingulate cortex left & right,
- 9 (HyThL) & 10 (HyThR): hypothalamus left & right.

#### 1) STATIC FC ANALYSIS

In order to study the resting-state FC in the different thermal conditions, the post-treated time course of the cerebral blood volume (CBV) signal of the N=10 ROIs (Figure 1 C, D) were extracted. Simply stated, the temporal signal during the calm periods (without motion artefacts) was extracted from each ROI. The NxN Pearson correlation matrix of the ROI signals was computed over time. First, each correlation coefficient of the correlation matrix was individually compared between pairs of conditions. Since the measurements were not always independent (because of intra and inter group comparison, and of repeated measurements on some animals, see Supplementary Tables 1 and 3), these comparisons were made with linear mixed models having the animal as random effect factor, and the thermal condition as two-modality fixed effect factor, whose significance was tested. The correlation coefficients being Fisher-transformed, the parameter estimation was made using restricted maximum likelihood estimation, and the validity of the model was checked posteriori by testing the normality of its residuals with Shapiro-Wilk’s test. Then, in order to account for multiple testing (a NxN correlation matrix involves Nx(N-1)/2 = 45 correlation coefficients), we performed Benjamini-Hochberg’s adjustment for multiple comparisons on the p-values of significance of the thermal condition effect. A false discovery rate of 0.05 was adopted.

#### 2) DYNAMIC FC ANALYSIS

In order to study the dynamic behavior of FC, the above correlation matrices can be decomposed into the contribution of each time point, i.e. the co-fluctuation matrices [Esfahlani et al. 2020]. As a matter of fact, consider **x**_i_ = [x_i_(1) … x_i_(T)]^T^ the time series of the CBV signal of ROI n°i, and **x**_j_ = [x_j_(1) … x_j_(N)]^T^ that of ROI n°j. On one hand, the correlation coefficient between ROIs n°i and j can be computed by first z-scoring each **x**_i_ according to **z**_i_ = (**x**_i_ - m_i_)/s_i_, where m_i_ = 1/T ∑_t_ x_i_(t) and s_i_^2^ = 1/(n-1) ∑_t_ (**X**_i_ - m_i_)^2^ are the empiric mean and variance of the time-series over time, and then by computing r_ij_ = **z**_i_^T^ **Z**_j_/(T-1). Hence a single NxN correlation matrix with only Nx(N-1)/2 = 45 elements of interest since the matrix symmetric with unit diagonal. On the other hand, it is possible to consider the element-wise product of **z**_i_ and **z**_j_ as encoding the magnitude of the moment-to-moment co-fluctuations between ROIs n°i and j, and the 3D array of the NxN couples of **z**_i_ and **Z**_j_ as a time series of T co-fluctutation matrices of size NxN, each with only Nx(N+1)/2 = 55 elements of interest since the co-fluctuation matrices are symmetric.

To assess the repetitive nature of the dynamic characteristics of brain networks as a response to thermal inputs in an unsupervised fashion, we performed k-means clustering of the co-fluctuation matrices contributing to the static correlation matrices. The time series data from all 92 acquisitions on all animals were concatenated together to form a single time series of T = 84 499 time points for each ROI, yielding a 3D array of size 84 499 x 10 x 10.

Prior to k-means clustering, outliers were removed. To this end, the 84 499 x 84 499 L1 distance matrix between cofluctuation matrices considered as Nx(N+1)/2 vectors was computed, and the matrices with mean distance to the others larger than 3 were discarded, decreasing to T = 83 325 the number of co-fluctuation matrices to be clustered. K-means clustering was performed on the T remaining matrices considered as Nx(N+1)/2 vectors using the L1 distance with k=3, 5, 6 and 7 clusters: for each value of k, to avoid local minima, the algorithm was run 500 times with random initializations of centroid positions, the configuration minimizing the total sum of intra-cluster distances being retained. Finally, each time point was assigned to a cluster (or brain state), resulting in a dynamic characterization of FC patterns. For each of the 4 choices of k, the occurrence rates of all brain states were calculated for each animal in all conditions. Prior to the comparison of these occurrence rates across thermal conditions, we checked the homogeneity of the animal distribution across the brain states (see Supplementary Table-2). In order to establish the significance of the effect of the thermal condition on the occurrence rate of each state, linear mixed models were used for the same reasons as for the elements of the correlation matrices (intra and inter group comparison, varying numbers of repeated measurements). As before, the animal was considered as a random effect factor, and the thermal condition as a fixed effect factor, this time with several modalities (the constant and varying temperature conditions). When the latter could be considered significant with a type I error risk of 5%, two-by-two comparisons were performed, and the corresponding p-values were adjusted using Benjamini-Hochberg’s procedure, a false discovery rate of 0.05 being again adopted.

## RESULTS

### Changes in brain functional connectivity during exposure to sustained neutral / warm or cold floor surface

In order to decipher the FC alterations in brain networks, which are indicators of dynamic changes in the brain, we developed an experimental setup based on a previously established protocol on awake and freely moving mice (Rabut et al., 2020), but this time allowing the imaging during the application of varying floor temperatures (Figure 1).

In the first part, we investigated the potential changes of FC in cortical and hypothalamic areas located in the chosen imaging plane during 20 minutes of exposure to a constantly neutral (25°C), warm (35°C) or cold (15°C) floor. After removing motion artifacts and concatenating the cleaned signals, the time series were first analyzed in a stationary way, i.e. averaged over the recording sessions.

Exposure to constant floor temperature: 35°C (warm) and 15°C (cool) temperatures for 20 minutes were compared to 25°C (neutral temperature). We observed large and statistically significant differences of FC between these conditions, with the largest number of significant differences for the cold (15°C) floor, for which the FC of 19 couples of ROI were statistically significantly different (Figure 2 A, B, C). Twelve of these pairs concerned areas of the somato-motor network (SMN), which had a stronger connectivity at 15°C than at 25°C (Figure 2 D). A single ROI pair, concerning cingulate and hypothalamus, exhibited an increase in connectivity at 15°C. The seven other pairs of ROI concerned areas between a ROI of the SMN and the hypothalamus (Figure 2 D). Interestingly, in these pairs, the FC was altered in the opposite way: the FC was significantly decreased (Figure 2 H, I).

**Figure 2:**
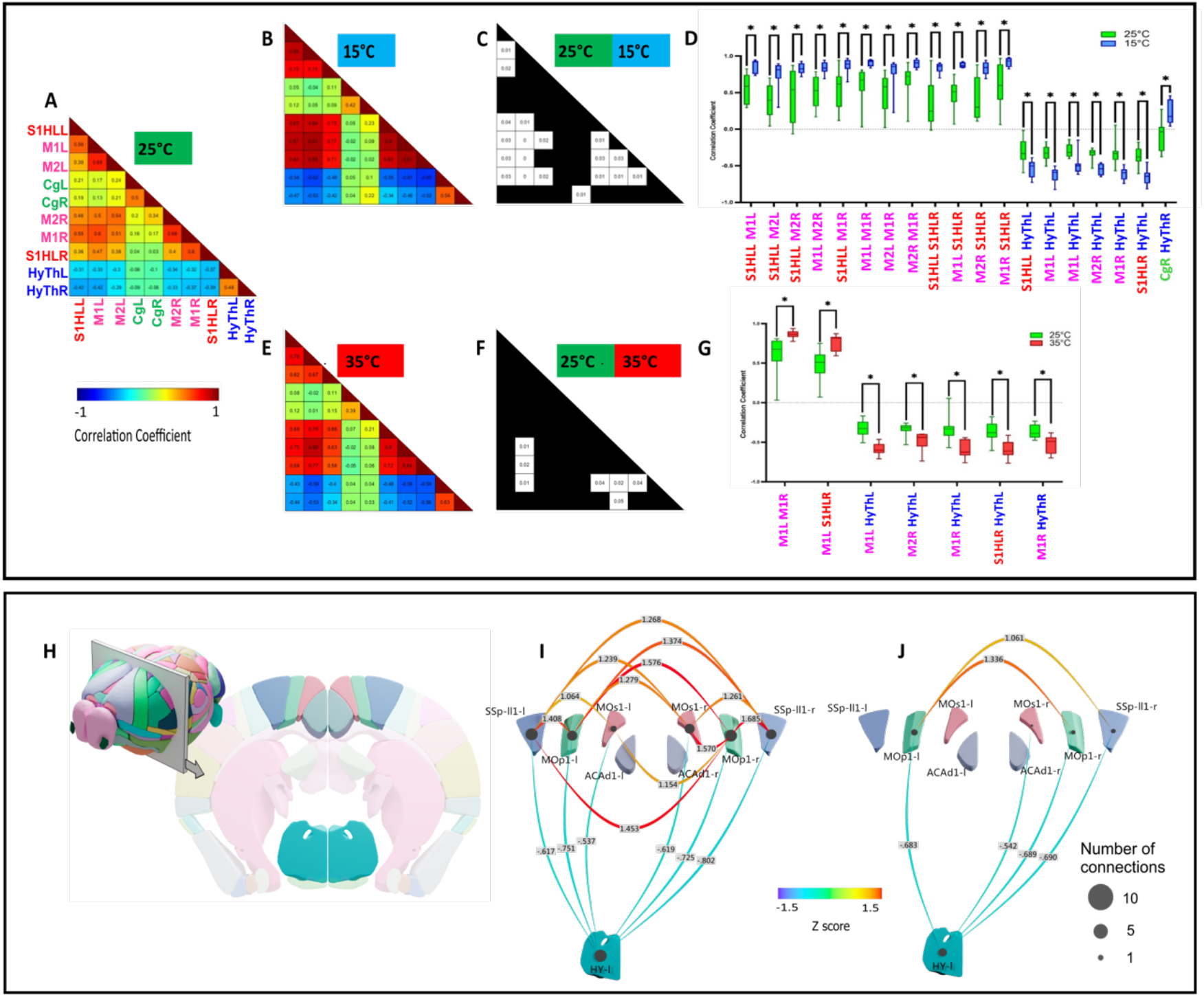
Functional connectivity comparison between constant 15°C, 25°C and 35°C using correlation matrices show striking differences between neutral (25°C) and cold (15°C) rather than neutral and warm (35°C). A) Average Pearson correlation matrix of N=8 imaging sessions at 25°C. (B, E) Same for imaging sessions at 15°C (N=8) and 35°C (N=8) respectively. (C, F) Significance matrix indicating the ROI pairs with significant differences between 25°C and 15°C and 35°C respectively. In these matrices, ROI pairs with significant alterations (adjusted p < 0.05) between any two conditions are indicated by white squares with the corresponding p-value. (D, G) Boxplot representation of each ROI pair with a significant FC alteration between the conditions 25°C vs 15°C, and 25°C vs 35°C. *p < 0.05, **p < 0.01 and ***p < 0.001 of linear mixed model analysis of the thermal condition effect, followed by Benjamini-Hochberg’s correction for multiple comparisons. (H-J) Summary representation in form of graphs of the statistically significant changes between 25°C and 15°C (I) and 25°C and 35°C (J) shown as Z-scores between these conditions. (H) Schematic of the imaging plane Bregma −0.34 mm in relation to the 3D whole mouse brain. I: Imaging plane with the regions studied (SSpll1-l/r - primary somatosensory cortex of hindlimb S1HL, MOp1-l/r - primary motor cortex M1, MOs1-l/r - secondary motor cortex M2, ACAd1-l/r cingulate Cg and HY l/r Hypothalamus).

Exposure to warm floors (35°C) induced mild changes of the brain FC (Figure 2 A, E, F). Only seven ROI pairs displayed a significant alteration of the FC. Two of these were pairs of the SMN, which displayed an increased FC (Figure 2 D). The five other ROI pairs concerned areas of the SMN and the hypothalamic nuclei (Figure 2*D*). They all displayed the opposite effect: an increased FC when the mice were submitted to a warmer floor (Figure 2 J).

To conclude, for both temperatures (15°C and 35°C), when the floor temperature is constant, the somato-motor regions display an increased connectivity, while the hypothalamic-somato-motor network connectivity decreases. By taking a closer look, it was observed that the connectivity between somato-motor network is higher at 15°C and 35°C than at 25°C, and that the hypothalamic-somato-motor network anti-correlation was also higher at 15°C and 35°C than at 25°C.

### Alterations of the stationary FC during fast dynamic changes of temperature

In order to decipher the neurobiological mechanism that takes place during dynamic changes of the temperature, we next compared the stationary FC during fast or slow, warm or cold ramps.

Warm Fast ramps did not lead to any significant FC difference compared to 25°C and 35°C conditions (Supplementary Figure 1). However, Down and Up Cold Fast ramps led to strong significant FC alterations compared to 25°C and 15°C conditions in numerous ROI pairs (Figure 3). The Cold Fast Ramps Down were the conditions that produced the largest number of statistically different ROIs, with 18 pairs of regions modified with respect to the neutral temperature (25°C), and 15 pairs when compared to the target temperature (15°C).

**Figure 3:**
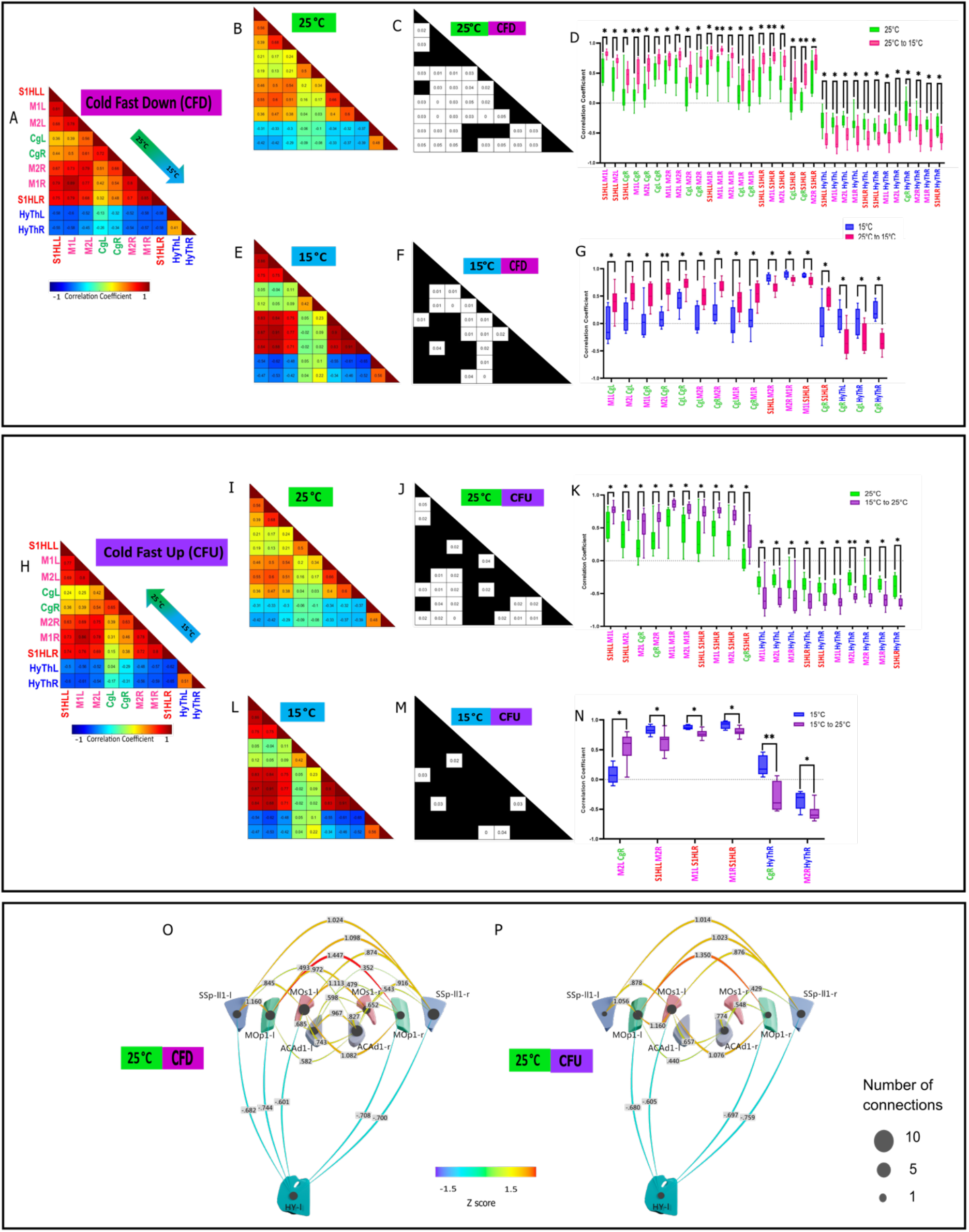
Functional Connectivity comparison between the Cool Fast Down (25°C to 15°C) and Cool Fast Up (15°C to 25°C) ramps on one hand, and 25°C and 15°C on the other hand. (A) Average Pearson correlation matrix of N=10 imaging sessions during Cool Fast Down. (B, E) Same for imaging sessions at constant 25°C and 15°C respectively. (C,F) Significance matrix indicating the ROI pairs with significant differences between 25°C and Cool Fast Down, and 15°C and Cool Fast Down respectively. (D,G) Boxplot representation of ROI pairs with a significant FC alteration between Cool Fast Down ramps, and 25°C and 15°C respectively. (H) Averaged Pearson correlation matrix of N=10 imaging sessions during Cool Fast Up. (I,L) Same for sessions at fixed 25°C and 15°C respectively. (J,M) Matrix indicating the ROI pairs with significant differences between Cool Fast Up ramps, and 25°C and 15°C respectively. (K,N) Boxplot representation of the ROI pairs with a significant FC alteration between Cool Fast Up and 25°C and 15°C respectively. *p < 0.05, **p < 0.01 and ***p < 0.001 of linear mixed model analysis of the thermal condition effect, followed by Benjamini-Hochberg’s correction for multiple comparisons. (O, P) Summary representation in form of graphs of the statistically significant differences shown in C and J, i.e. Z-scores between the conditions: CFD vs 25°C (O) and CFU vs 25°C (P).

When compared to 25°C (Figure 3 A, B, C), 5 pairs of regions involving the cingulate and SMN, and 7 couples within the SMN displayed an increased FC when the temperature was lowered (Figure 3 D). The last 6 ROI pairs concerned the hypothalamus and SMN or the cingulate (Figure 3 D). They all had an initial anti-correlation at 25°C, which became even stronger when the floor became colder (Figure 3 D, O).

The comparison to constant 15°C floor temperature revealed 15 pairs of regions with significantly modified FC (Figure 3 A, E, F). Among them, 13 encompassed the SMN, where the FC increased when the floor temperature decreased (Figure 3 G). Two (intra-motor network and motor-S1HL) had the opposite behavior: slight but statistically significantly decreased FC (Figure 3 G). Finally, as shown in panel D of Figure 3, networks between the hypothalamus and the cingulate showed anticorrelations when the floor was getting colder (Figure 3 G).

Overall, these results suggest that a fast decrease in temperature is associated with a strengthening of the correlation of the somato-motor network, and a weakening of the link between the hypothalamus and the somato-motor network (Figure 3 O).

The analysis of the Cold Fast ramp Up (Figure 3 H-N) showed a similar effect in a smaller number of regions. Compared to the target temperature (25°C), 5 pairs of regions within the SMN showed an increased FC during the ramp (Figure 3 J, K). Seven couples involving the hypothalamus and areas of the SMN displayed an increased anti-correlation of their FC (Figure 3 J, K). The comparison of this Fast-Cold Up ramp to the initial temperature (15°C) revealed that only 6 pairs of regions (Figure 3 M, N) differed significantly (4 within the SMN and 2 between the hypothalamus and the cingulate and motor cortex).

The striking difference between the two Cold Fast ramps (Down and Up) was the large number of subnetworks within the SM-cingulate networks in which the FC was reinforced. In both cases (Up and Down), the hypothalamus-SM-cingulate networks which are slightly anti-correlated in neutral conditions, display a stronger anti-correlation during these ramps (Figure 3 O-P). These results highlight the complex inter-regional FC between the cingulate, somatosensory and motor cortices and hypothalamus during exposure to cold.

### Effect of the rate of temperature change

In order to further understand how changes of floor temperature are processed centrally, we also investigated the effect of the temperature change at a smaller rate of 0.5°C/min (20 min duration), called Cold Slow/ Warm Up/Down, that we compared with the fast ramp initially used (1°C/min, 10 min duration). We observe that, as hypothesized, these contrasting conditions produce strikingly different effects. As a matter of fact, the Warm Slow ramps only lead to slight FC alterations (Supplementary Figure 2).

Only 8 pairs of ROI were significantly different between the Cold Slow Down ramp and constant 25°C (Figure 4 A, B, C), which is roughly half of what we observed with the fast equivalent of this ramp. As observed with the fast-cold ramp, the modified ROI pairs concerned the SMN (in which the FC was reinforced), and the SMN and hypothalamus, where the anti-correlation was also reinforced (Figure 4 D).

**Figure 4:**
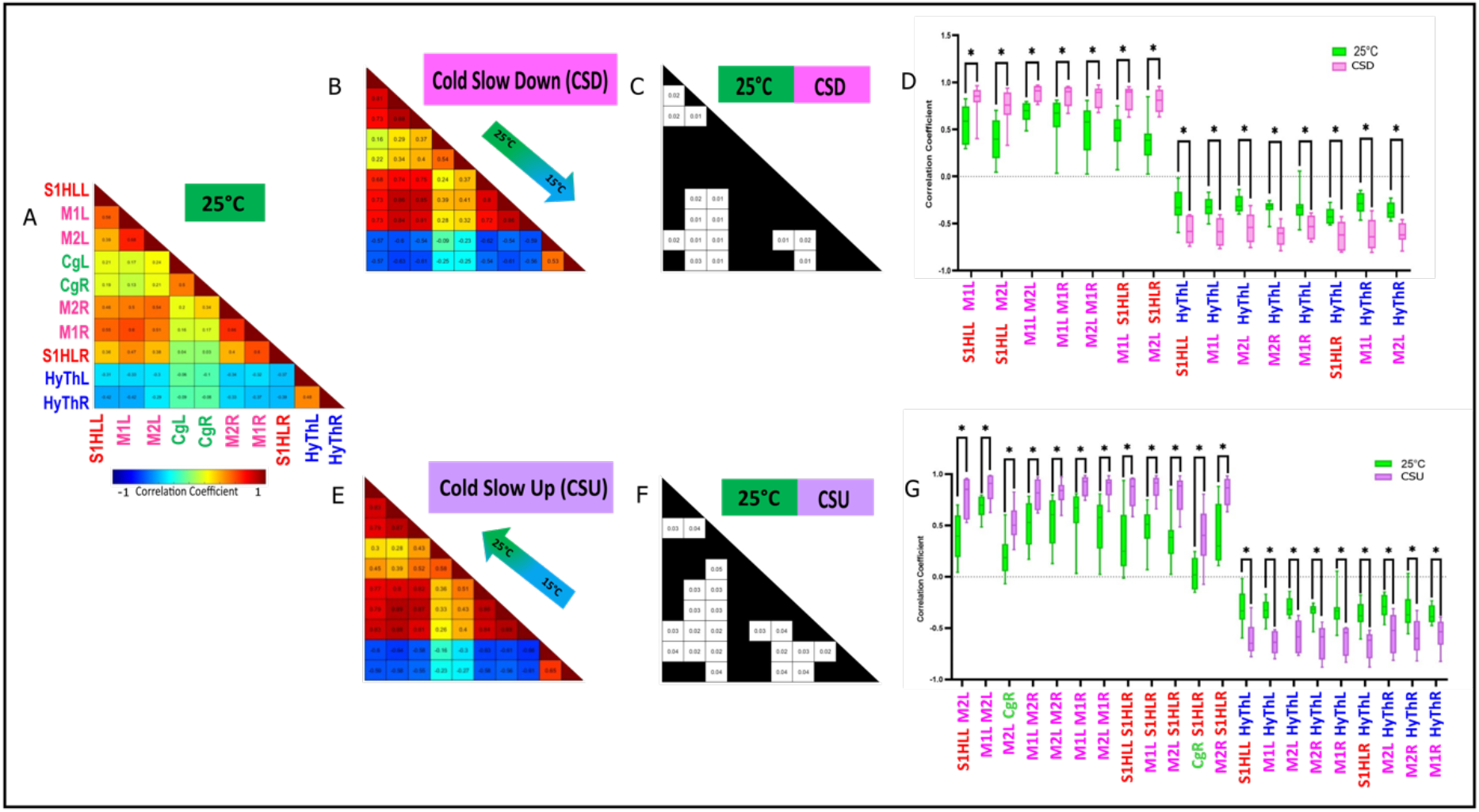
Functional Connectivity comparison between the 25°C condition, and the Cool Slow Down (25°C to 15°C) and Cool Up (15°C to 25°C) ramps. (A) Average Pearson correlation matrix of N=8 imaging sessions at 25°C. (B,E) Same for the Cool Slow Down (N=8) and Cool Slow Up (N=6) imaging sessions. (C, F) Significativity matrix indicating the ROI pairs with significant differences between 25°C with Cool Slow Down and Cool Slow Up ramps respectively. (D, G) Boxplot representation of the ROI pairs with significant FC alterations between 25°C and Cool Slow Down and Cool Slow Up ramps respectively. *p < 0.05, **p < 0.01 and ***p < 0.001 of linear mixed model analysis of the thermal condition effect, followed by Benjamini-Hochberg’s correction for multiple comparisons.

As for the Cold Up ramp, in contrast with the aforementioned Cold Slow Down ramp and very similarly to the fast ramps, it displayed a large number of pairs of ROI modified. Most of them were within the SMN (12 out of 21 (Figure 4 F,G) with a strengthening of the FC between them. The anti-correlation between the hypothalamus and the SMN was significantly reinforced here (Figure 4 G), as observed in (Figure 3 D).

In conclusion, the analysis of the effect of the rate of temperature change shows that the larger the rate, the larger the number of ROI with statistically significant changes, especially in the Cold Down ramps. Another important difference is the lack of significant differences between these slow ramps and the constant low temperature of 15°C, suggesting that when the ramp is slow, the overall central processing does not differentiate well between a cold floor and one that is slowly getting cooler.

### Dynamic FC

The dynamic nature of the stimuli applied in this study and the rapid changes of brain states occurring in awake and conscious animals naturally led to the analysis of the dynamic connectivity. We predicted that the FC changes during thermal variations might be even stronger than the differences between warm and cold conditions.

In a previous study, we established that fUS imaging can robustly identify brain states extracted from k-means clustering of steady state fUS data in anesthetized rats (Rahal et al., 2020). In the current study, using a similar approach, but adapted to data discontinuity generated by artifact removal, we could robustly identify 5 consistent brain states using k-means with K=5, 6 or 7 (Figure 5 A-C). As a matter of fact, whatever the value of k, the 5 states are not only reproducible, but also display a stable percentage of occurrence across time. Finally, the statistical effects observed in our experimental groups were consistently obtained (with mild variation for state 6).

**Figure 5:**
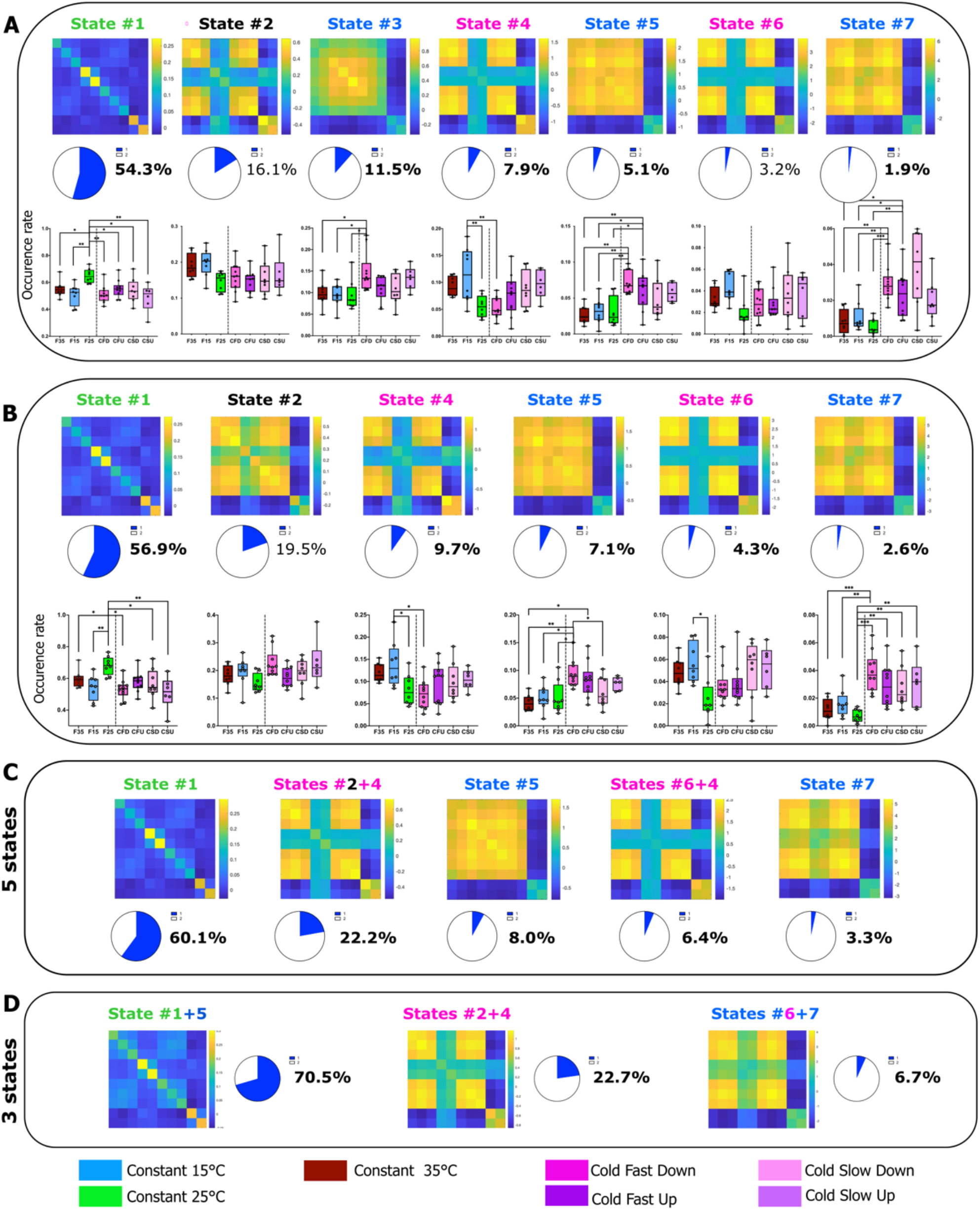
Dynamic functional connectivity analysis of thermal coding using k-means clustering for either k=7, 6, 5 or 3. (A) 7 brain state matrices (cofluctuation matrices) and (B) 6 brain state matrices were ordered according to their decreasing level of connectivity with their probability of occurrence in cold and neutral conditions, and significance of the thermal condition effect using linear mixed models. *p < 0.05, **p < 0.01 and ***p < 0.001 of linear mixed model analysis of the thermal condition effect. Brain States #3, #5 and #7 show dichotomy between SM/Cg and hypothalamus and occur more often in Cold Fast Down ramps, whereas states #4 and #6 predominantly occur during sustained cold exposure and display a decreased connectivity between Cg and other regions. The weakly correlated state #1 is present in all thermal conditions, but its significantly more frequent in the 25°C condition. (C-D) k-means clustering of brain states into 5 and 3 states, respectively.

First, the warm ramps did not produce any significant difference in FC compared to the constant temperatures (Supplementary figure 3). Among the brain states (identified with the k=7 algorithm and labeled 1 to 7, see Figure 5A), the strengths of the connections and the statistical results suggest four groups of states. The first group comprises state #1 only, which is by far the most frequent state (54-60% of the time, Figure 5A-C), has weak connections, and is consistently more present at neutral temperature (constant 25°C) compared to all other experimental conditions (Figure 5A-B).

The second group is made up of states #3, #5 and #7 which are present 10-17% of the time are characterized by a common pattern of connectivity in the SM-cingulate (SM-Cg) cortex and an anti-correlation between the SM and the hypothalamic regions. Interestingly, these three states were significantly more frequent during the CFD ramp, compared to all constant temperatures.

The third group consisted of states #4 and #6. In addition to the dichotomy between the SM and hypothalamic networks, these states (present in 10-13% of the time)present are characterized by a low correlation between the cingulate and all other areas. Very interestingly, states #4 and #6 were consistently more frequent in one condition: the low constant temperature (15°C), suggesting a specific involvement of the cingulate cortex during this prolonged (20 minutes) exposure to cold.

Finally, the last group consisted in the single state 2, which was present 16-19% of the time and that was statistically equally represented in all experimental conditions.

## DISCUSSION

The peripheral and central neurophysiological mechanisms involved in thermal sensing have been at the heart of many studies since the last century. In the present study, we aimed at deciphering the interplay between the brain regions that are already shown to be crucial for thermal processing. Taking advantage of the sensitivity and versatility of fUS Doppler imaging in awake freely moving animals, we analyzed the stationary and dynamic changes of FC, an indirect marker of brain network functionality and dynamic characteristics. Despite the technical limitations of our approach, our study brings forward robust information regarding the complexity of interplay between the somato-motor and the hypothalamic brain networks.

### FC: A strong marker of strength of interaction between brain regions

The analysis of intrinsic brain activity has led to the definition of resting brain state networks. Since then, the field of neuroscience has been widely studying these infra-slow oscillations, called functional connectivity (FC), that characterize the spatiotemporal organization and function of large-scale brain networks (Preti et al., 2017)(Preti et al., 2017). Interestingly, FC can be studied using very different approaches, such as fMRI, electrophysiology, MEG, fiber photometry, optical imaging and fUS (Pais-Roldán et al., 2021). Although the classical studies focused on resting state FC, task or behavior related FC studies using fMRI has also been increasing in numbers in the recent years (Barch et al., 2013; Cole et al., 2014; Di and Biswal, 2019). Although the mice are not subjected to a task in our study, they are physiologically and behaviorally responding to the thermal stimulation which is either cold, warm or neutral. fUS has been previously shown to be able to measure FC in anesthetized rats (Osmanski et al., 2014b), task-evoked changes in FC (Ferrier et al., 2020), pharmacological studies in rodents (Rabut et al., 2020), but also altered FC in preterm babies (Baranger et al., 2021). Unlike the conventional analysis of time averaged FC that provides quantitative information on the correlation (or anti-correlation) between pairs of regions of interest (ROI) in the steady-state, there have been tremendous advancements in the study of the dynamic nature of FC (Hutchison et al., 2013; Preti et al., 2017). The temporal evolution of FC can reveal how FC reshapes according to the physiological (Tarun et al., 2021) or behavioral changes, at rest, or during a task, or in case of neurodegenerative or neuro-psychiatric illnesses (Tian et al., 2018; Barttfeld et al., 2019; Demertzi et al., 2019; Gu et al., 2020).

### Warm conditions do not elicit FC alterations

The temperature conditions in the present study were carefully designed as not to induce any discomfort in the mice during awake imaging. Although 25°C falls in a slightly low temperature range, adaptation takes place before the beginning of the experiment and can be considered neutral. Thermal psychophysics studies have shown that, when the skin is adapted to temperature values ranging from ~30 to ~34°C, neither warm nor cool sensations are experienced (Filingeri, 2016). Furthermore, the transition from ambient to high temperature between 32-39°C and 26-34°C activates the TRPV3 and TRPV4 channels respectively. The TRPM2 channel is activated at approximately 35°C by sensing environmental temperatures (Tan and McNaughton, 2016). Therefore, it is likely that a temperature of 35°C would not either have evoked a significantly different response from that observed at neutral temperature.

### Dichotomy between somato-motor-cingulate and hypothalamic networks during exposure to cold

In neutral conditions, the stationary FC indicated that, as previously demonstrated in fMRI in human (Zeng et al., 2012) and rodents (see for review (Pais-Roldán et al., 2021)), the SMN has a strong positive interhemispheric connectivity, that was shown previously to be due to a strong concomitant interhemispheric neuronal activity. Our study demonstrates a reinforcement in the somato-motor FC during cold sensing, mostly in the Cold Fast ramps. We hypothesize that such a FC increase is an indirect readout of the sensory discriminative aspect of cold sensing.

In contrast with this, the SM-hypothalamic network shows a decreased FC during cold sensing, suggesting a dichotomy between these two networks. A similar dramatic opposing effect was also observed when analyzing the dynamic FC. Indeed, the second type of dynamic modes reproducibly obtained in our analysis were the states 3, 5 and 7, which are characterized by a dichotomy between a strongly correlated signal within the SM-Cg network and an anti-correlation in the SM-hypothalamic network. These three modes, which account for approximately 10-17% of the time, have a higher occurrence during the Cold Fast Down ramp. The significant differences with other conditions, including the Cold Slow Down ramps and the constant cold condition (15°C), suggest that this dichotomy is a specific feature of cold sensing.

### Weakly connected dynamic mode in thermal sensing

By measuring the temporal fluctuations in FC, we were able to identify the dynamic patterns using k-means clustering. The results were robust for k=5, 6, 7, the nature of the states and the statistical effects observed being similar whatever k. These states, that we named 1 to 7 (clustering with k=7), encompass different neurobiological meaning and role.

The first group of states consists of state #1 alone. As previously described in monkeys (Barttfeld et al., 2019) and human subjects (Demertzi et al., 2019) using the same type of analysis but in fMRI, the most frequent dynamic state (50-60% of the recording time) is a mode characterized by weak strengths of connection. Our current understanding of this state is that it is associated with low-level cognitive functions (Barttfeld et al., 2019; Demertzi et al., 2019). In our study, while present in the majority of all experimental groups, it is statistically more frequent when the animals were exposed to the neutral temperature of 25°C. This result reinforces previous suggestions of low FC, apart from the default mode network during resting periods.

### A secondary pathway involving hypothalamus

The hypothalamus was anti-correlated to the somato-motor network in all conditions and the strength of this anti-correlation was the highest during cold sensing. This dramatic effect was revealed in both stationary and dynamic FC analysis. Although the preoptic anterior hypothalamus (POAH) has been linked to thermoregulatory behavior (Ishiwata et al., 2002; Wang et al., 2019), the secondary thermosensory pathway from dorsal horn to the lateral parabrachial nucleus (LPB) and then to the preoptic area (POA) of the hypothalamus has been put forward by (Yahiro et al., 2017). Using selective lesions of either thalamic nuclei or the POA and the LPB, they unraveled an important role of these nuclei in thermoregulatory behaviors (Yahiro et al., 2017). Our hypothesis is that the changes observed in the hypothalamus in our study are linked to this role in thermoregulation.

Only a small number of FC studies in rodents are documenting networks involving hypothalamic nuclei. In the rare cases doing so, they define them as the ‘ventral midbrain’ (Liska et al., 2015) or the ‘basal ganglia-hypothalamus’ (Becerra et al., 2011). In agreement with our observations, these networks are different from the sensory network (Hutchison et al., 2010; Zerbi et al., 2015). If we postulate that this anti-correlation is due to a decrease in neuronal activity, this second structure would have an inhibitory action on hypothalamic neurons. However, this anti-correlation can also be due to a time lag between signals of the SMN and the hypothalamic. In this last case, the hypothesized structure(s) would only change the temporal shift, without affecting the intensity of neuronal activity.

### Sustained cold and cold ramps are encoded differentially in the cingulate corteX

As previously described by us and by others (Osmanski et al., 2014b; Liska et al., 2015; Zerbi et al., 2015), correlation between regions of the SMN and the cingulate cortex was much weaker, as it belongs to the default mode network (Liska et al., 2015). During Fast Cooling, interhemispheric connectivity within the cingulate cortices increased significantly, as well as the connectivity between the SM and the cingulate cortices. When the cold ramp was slow, however, the FC between these regions was not affected. The thermal psychophysics associated with rapid temperature changes and the already established role of the cingulate cortices in affective responses to unpleasant or nociceptive thermal sensations (Craig et al., 1996; Becerra et al., 1999; Brooks et al., 2002; Derbyshire et al., 2002) has led us to consider that the concomitant FC increase between SM and cingulate cortex suggests a common hub between the two.

In the dynamic brain states #4 and #6, on the other hand, the cingulate cortex is differently connected to the other networks during static exposure to a lower temperature (15°C). The decrease in connectivity with the somato-motor network in this state seems to be a characteristic of sustained exposure to cold, and this suggests the differential role of cingulate in sensing persistent cold sensations and cold ramps.

## Authors contribution statement

SP, MT and HK designed the experimental paradigm.

SC, AB and HK were involved in the awake functional ultrasound imaging and thermal experimental setup.

HK and SP wrote the manuscript.

HK performed the experiments and analyzed the ultrasound data.

MT, TD and JS supervised the signal processing of the ultrasound data.

IR and YL performed the statistical analysis.

AD was involved in the signal processing.

SP, MT, JS, HK and IR were involved in the interpretation of the data and wrote some parts of the manuscript.

## Acknowledgments

The authors wish to thank Nathalie Ialy-Radio for animal husbandry and the CNRS, INSERM and ESPCI for their financial support. This work was supported by a funding from the European Union’s Horizon 2020 research and innovation program under the Marie Skłodowska-Curie grant agreement No 754387 (PhD fellowship Miss Koorliyil) and from the Agence Nationale de la recherche (Project ‘PINCH’, 18-CE37-0005-01). In addition, this work was supported by the Chair in Biomedical Imaging of the AXA Research Fund and the European Research Council (ERC) Advanced Grant FUSIMAGINE.

## Supplementary Figures

**Supplementary Figure 1:**
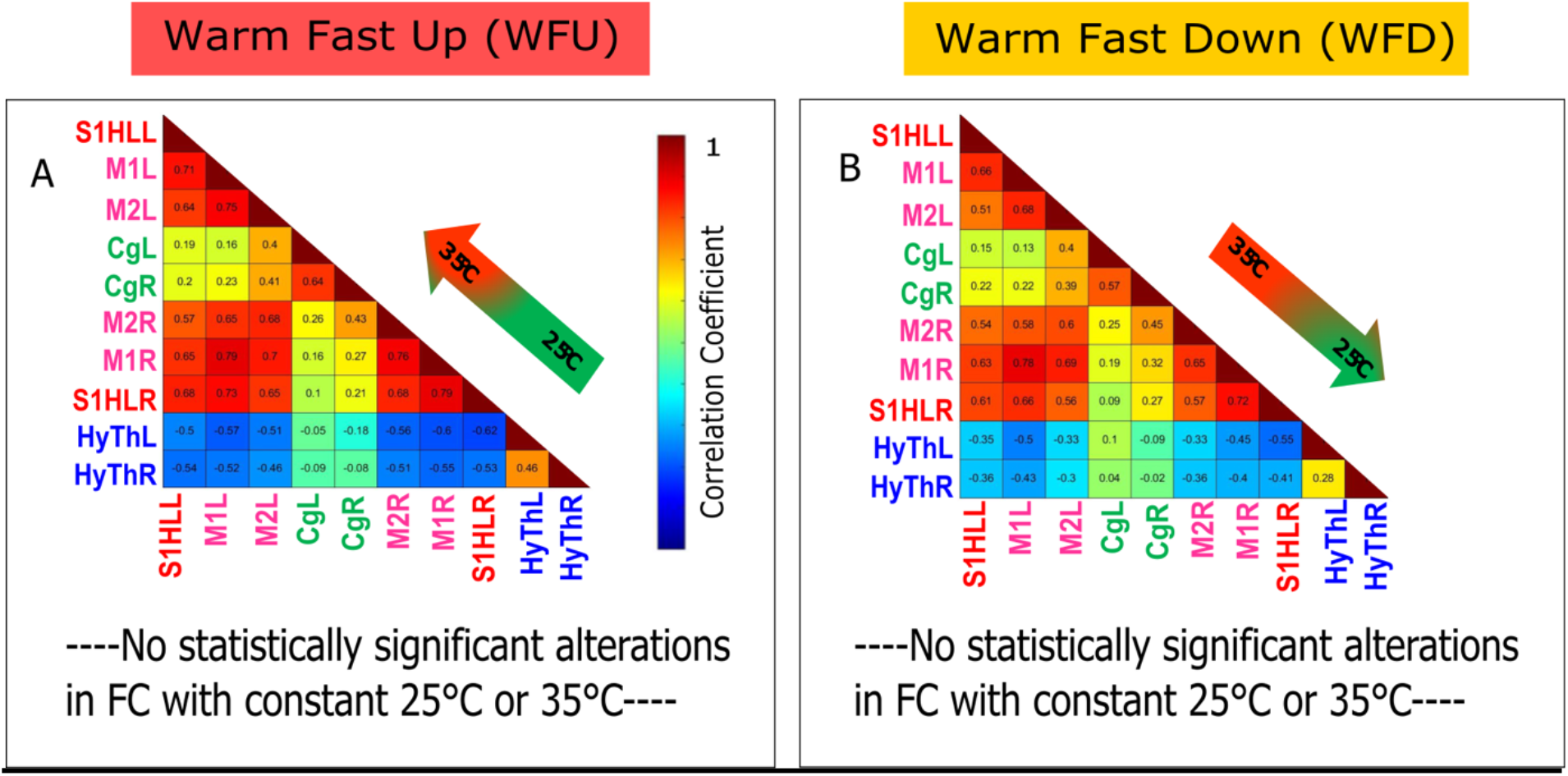
No changes in functional connectivity are observed between Warm Fast Up, Down ramps and either 35°C or 25°C conditions using correlation matrices. (A, B) Averaged Pearson correlation matrices of WFU (N=8) and WFD (N=8) imaging sessions. (C, E). Average Pearson correlation matrix of N=8 imaging sessions at 25°C. No significant FC alteration was observed.

**Supplementary Figure 2:**
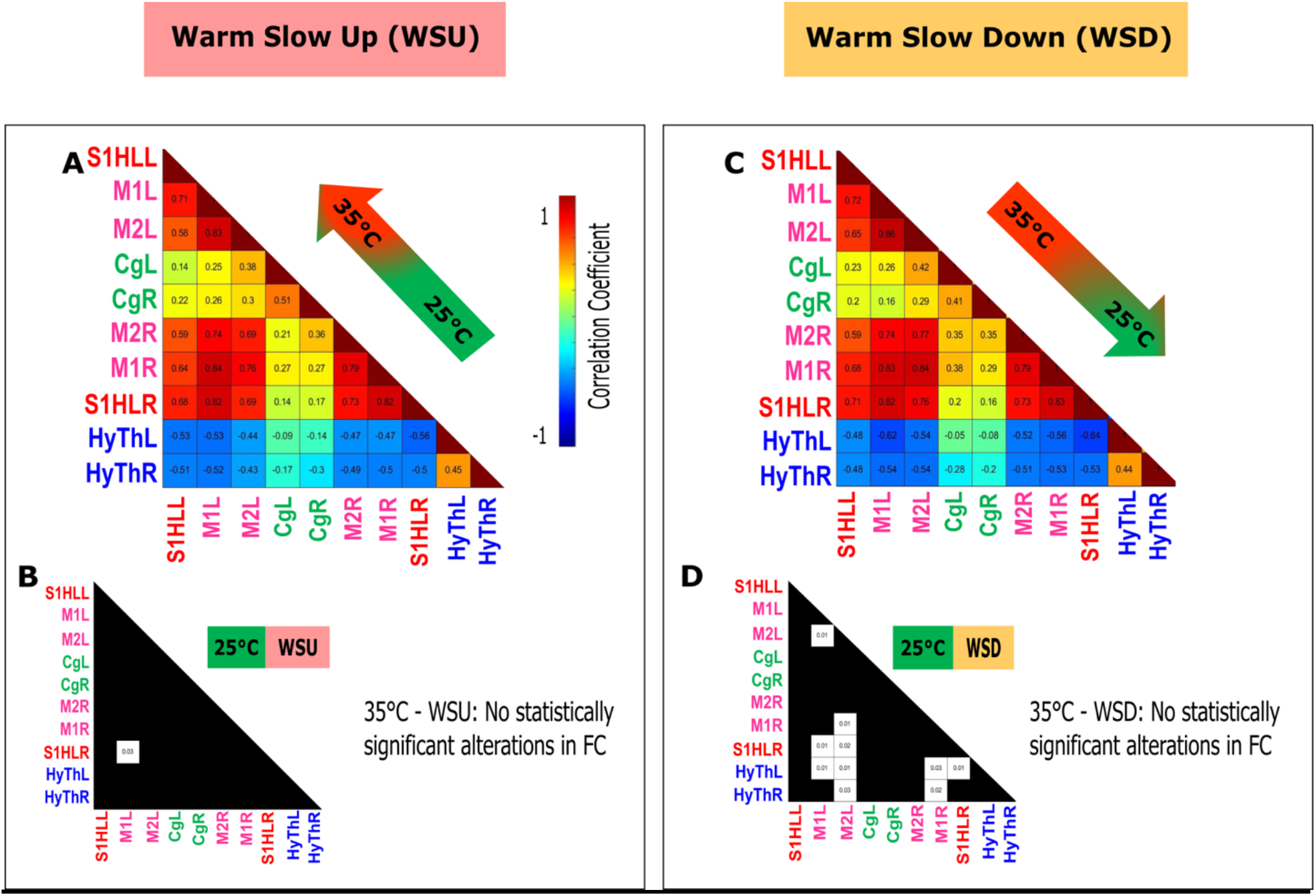
Functional connectivity comparison of Warm, Slow Up (A, B) and Slow Down ramps (C, D) with 35°C and 25°C conditions using correlation matrices show limited changes. (A, C) Average Pearson correlation matrices during WSU (N=8) and WSD (N=8) imaging sessions. (B, D) Matrix indicating the ROI pairs with significant differences between WSU and 25°C and WSD and 25°C respectively. No statistical differences were observed between the constant 35°C temperature and the conditions WSU and WSD.

**Supplementary figure 3:**
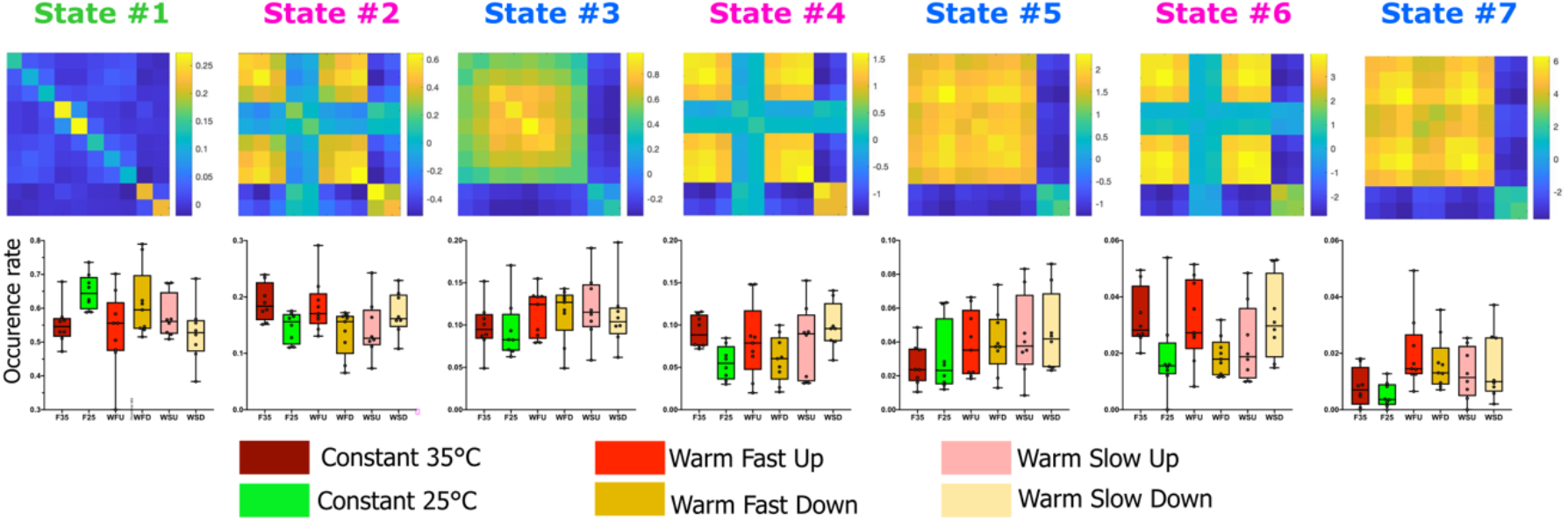
Lack of statistical difference in the occurrence rate of the various dynamic functional connectivity states in warm experiments (constant and ramps). As presented in figure 5, dynamic functional connectivity was analyzed using k-means clustering with k= 7, which are ordered from 1 to 7 in the order of decreasing occurrence rate. The frequency of occurrence was calculated for each brain state for all thermal conditions. No significant difference could be established using a linear mixed model analysis of the thermal condition.

**Supplementary Table 1:** Reporting of included animals and their number of imaging sessions. Table presenting the mice (M) included in each part of the study and the number of sessions kept for each one of them. The numbers indicate how many sessions of each mouse was kept. The number following ‘M’ is the mouse number. Ex/ M401: Mouse #401. Grey boxes: due to artifacts, the signals obtained for this recording was too noisy and had therefore to be discarded (see materials and methods).

| Nb Mouse | Constant 25°C | Constant 15°C | Constant 35°C |
| --- | --- | --- | --- |
| M401 | 1 | 1 | 1 |
| M402 | 1 | 2 | 1 |
| M403 | 2 | 2 | 2 |
| M404 | 1 | 2 | 2 |
| M405 |  |  | 1 |
| M406 | 1 | 1 | 1 |
| M302 | 1 |  |  |
| M304 | 1 |  |  |
| **Total sessions** | **8** | **8** | **8** |

II. Number of imaging sessions kept in Ramp experiments
|  | Cold Fast Down | Cold Fast Up | Cold Slow Down | Cold Slow Up |
| --- | --- | --- | --- | --- |
| M501 | 3 | 3 | 3 | 3 |
| M502 | 3 | 3 |  |  |
| M503 | 2 | 2 | 2 | 1 |
| M504 |  |  |  |  |
| M505 | 2 | 2 | 3 | 2 |
| M506 |  |  |  |  |
| **Total sessions** | **10** | **10** | **8** | **6** |

|  | Warm Fast Down | Warm Fast Up | Warm Slow Down | Warm Slow Up |
| --- | --- | --- | --- | --- |
| M501 | 3 | 3 | 3 | 3 |
| M502 | 2 | 2 |  |  |
| M503 | 2 | 2 | 2 | 2 |
| M504 |  |  |  |  |
| M505 | 2 | 2 | 3 | 3 |
| M506 |  |  |  |  |
| **Total sessions** | **9** | **9** | **8** | **8** |

**Supplementary Table 2:** Distribution of the different animals in the various brain states (related to Figure 5)

6 states
| Mouse # | State1 | State2 | State3 | State4 | State5 | state6 |
| --- | --- | --- | --- | --- | --- | --- |
| M302 | 0,010779897 | 0,011388942 | 0,005062993 | 0,01758794 | 0,003673354 | 0,006692694 |
| M304 | 0,013001708 | 0,007189667 | 0,005180737 | 0,004638578 | 0,003108223 | 0,000557724 |
| M401 | 0,031599087 | 0,030412929 | 0,029671494 | 0,03614225 | 0,02627861 | 0,029001673 |
| M402 | 0,043263593 | 0,044983139 | 0,041563641 | 0,031117124 | 0,029669398 | 0,023424428 |
| M403 | 0,070460203 | 0,045746644 | 0,075827152 | 0,02995748 | 0,06301215 | 0,021751255 |
| M404 | 0,059639161 | 0,069160781 | 0,090074179 | 0,0618477 | 0,097485165 | 0,045175683 |
| M405 | 0,016889876 | 0,016351721 | 0,016719651 | 0,012756088 | 0,014693416 | 0,01115449 |
| M406 | 0,037379909 | 0,035948336 | 0,055104203 | 0,021453421 | 0,058208533 | 0,012269939 |
| M501 | 0,268674525 | 0,286505058 | 0,233721889 | 0,302860456 | 0,252896298 | 0,321807027 |
| M502 | 0,101524409 | 0,096837819 | 0,055810668 | 0,118283726 | 0,062447019 | 0,149470162 |
| M503 | 0,129153038 | 0,135903798 | 0,14600259 | 0,141669888 | 0,149194688 | 0,158951478 |
| M505 | 0,217634594 | 0,219571165 | 0,245260803 | 0,22168535 | 0,239333145 | 0,219743447 |

7 states
| Mouse # | State1 | State2 | State3 | State4 | State5 | state6 | state7 |
| --- | --- | --- | --- | --- | --- | --- | --- |
| M302 | 0,011027972 | 0,007068178 | 0,015777611 | 0,005087949 | 0,014567035 | 0,004744958 | 0,008227375 |
| M304 | 0,013203115 | 0,006700044 | 0,008264463 | 0,004070359 | 0,003237119 | 0,003954132 | 0 |
| M401 | 0,03175708 | 0,031733176 | 0,032628528 | 0,025003634 | 0,035878069 | 0,023329379 | 0,029169783 |
| M402 | 0,043307087 | 0,049403622 | 0,037458409 | 0,038377671 | 0,024817912 | 0,030446817 | 0,020194465 |
| M403 | 0,06977857 | 0,064276248 | 0,038639047 | 0,072684983 | 0,020231994 | 0,06405694 | 0,020194465 |
| M404 | 0,057837038 | 0,083713739 | 0,059568531 | 0,089693269 | 0,058807661 | 0,101621194 | 0,04038893 |
| M405 | 0,016661591 | 0,017449566 | 0,016743587 | 0,014973107 | 0,012948476 | 0,014234875 | 0,008975318 |
| M406 | 0,036933919 | 0,046016787 | 0,026188687 | 0,056548917 | 0,019422714 | 0,060498221 | 0,005983545 |
| M501 | 0,269913429 | 0,269032543 | 0,277449823 | 0,251926152 | 0,31211222 | 0,235666271 | 0,33881825 |
| M502 | 0,10442859 | 0,076719187 | 0,103788773 | 0,053496148 | 0,132452118 | 0,061289047 | 0,155572177 |
| M503 | 0,129399226 | 0,130245914 | 0,146077063 | 0,146823666 | 0,139465875 | 0,158165283 | 0,145100972 |
| M505 | 0,215752382 | 0,217640995 | 0,237415477 | 0,241314144 | 0,226058808 | 0,241992883 | 0,22737472 |

**Supplementary Table 3:**
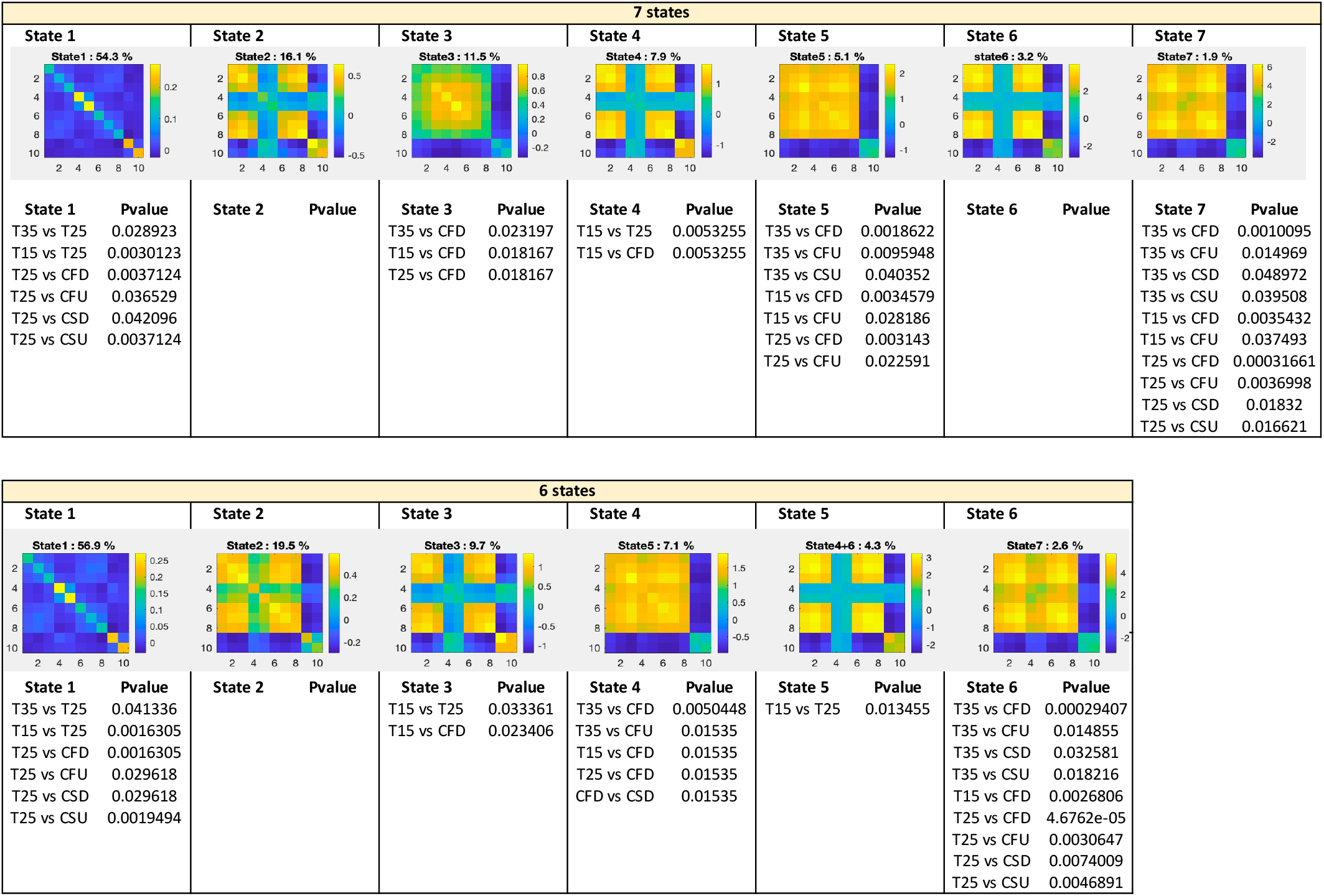
Statistical results of the mixed linear model.

| 7 states |  |  |  |  |  |  |
| --- | --- | --- | --- | --- | --- | --- |
| State 1 | State 2 | State 3 | State 4 | State 5 | State 6 | State 7 |
| **State 1** **Pvalue**<br>T35 vs T25 0.028923<br>T15 vs T25 0.0030123<br>T25 vs CFD 0.0037124<br>T25 vs CFU 0.036529<br>T25 vs CSD 0.042096<br>T25 vs CSU 0.0037124 | **State 2** **Pvalue** | **State 3** **Pvalue**<br>T35 vs CFD 0.023197<br>T15 vs CFD 0.018167<br>T25 vs CFD 0.018167 | **State 4** **Pvalue**<br>T15 vs T25 0.0053255<br>T15 vs CFD 0.0053255 | **State 5** **Pvalue**<br>T35 vs CFD 0.0018622<br>T35 vs CFU 0.0095948<br>T35 vs CSU 0.040352<br>T15 vs CFD 0.0034579<br>T15 vs CFU 0.028186<br>T25 vs CFD 0.003143<br>T25 vs CFU 0.022591 | **State 6** **Pvalue** | **State 7** **Pvalue**<br>T35 vs CFD 0.0010095<br>T35 vs CFU 0.014969<br>T35 vs CSD 0.048972<br>T35 vs CSU 0.039508<br>T15 vs CFD 0.0035432<br>T15 vs CFU 0.037493<br>T25 vs CFD 0.00031661<br>T25 vs CFU 0.0036998<br>T25 vs CSD 0.01832<br>T25 vs CSU 0.016621 |

| 6 states |  |  |  |  |  |
| --- | --- | --- | --- | --- | --- |
| State 1 | State 2 | State 3 | State 4 | State 5 | State 6 |
| **State 1** **Pvalue**<br>T35 vs T25 0.041336<br>T15 vs T25 0.0016305<br>T25 vs CFD 0.0016305<br>T25 vs CFU 0.029618<br>T25 vs CSD 0.029618<br>T25 vs CSU 0.0019494 | **State 2** **Pvalue** | **State 3** **Pvalue**<br>T15 vs T25 0.033361<br>T15 vs CFD 0.023406 | **State 4** **Pvalue**<br>T35 vs CFD 0.0050448<br>T35 vs CFU 0.01535<br>T15 vs CFD 0.01535<br>T25 vs CFD 0.01535<br>CFD vs CSD 0.01535 | **State 5** **Pvalue**<br>T15 vs T25 0.013455 | **State 6** **Pvalue**<br>T35 vs CFD 0.00029407<br>T35 vs CFU 0.014855<br>T35 vs CSD 0.032581<br>T35 vs CSU 0.018216<br>T15 vs CFD 0.0026806<br>T25 vs CFD 4.6762e-05<br>T25 vs CFU 0.0030647<br>T25 vs CSD 0.0074009<br>T25 vs CSU 0.0046891 |

## Data availability statement

Source data are available on the repository website Dryad using the following link: https://doi.org/10.5061/dryad.mkkwh713t.

## Code availability statement

Classical codes used to generate the results are available in the following depository: https://doi.org/10.5061/dryad.mkkwh713t. Custom codes used for the analysis of fUS data used in this study are protected by INSERM.

